# Distinguishing imported cases from locally acquired cases within a geographically limited genomic sample of an infectious disease

**DOI:** 10.1101/2022.07.15.500228

**Authors:** Xavier Didelot, David Helekal, Michelle Kendall, Paolo Ribeca

## Abstract

The ability to distinguish imported cases from locally acquired cases has important consequences for the selection of public health control strategies. Genomic data can be useful for this, for example using a phylogeographic analysis in which genomic data from multiple locations is compared to determine likely migration events between locations. However, these methods typically require good samples of genomes from all locations, which is rarely available. Here we propose an alternative approach that only uses genomic data from a location of interest. By comparing each new case with previous cases from the same location we are able to detect imported cases, as they have a different genealogical distribution than that of locally acquired cases. We show that, when variations in the size of the local population are accounted for, our method has good sensitivity and excellent specificity for the detection of imports. We applied our method to data simulated under the structured coalescent model and demonstrate relatively good performance even when the local population has the same size as the external population. Finally, we applied our method to several recent genomic datasets from both bacterial and viral pathogens, and show that it can, in a matter of seconds or minutes, deliver important insights on the number of imports to a geographically limited sample of a pathogen population.

## INTRODUCTION

Many infectious disease pathogens spread mostly within multiple geographical locations, for example countries, and are also occasionally imported from one location to another. When pathogen genetic data is available from several locations, a phylogeographic approach can be used to infer past migrations between countries (Lemey et al. 2009; Bloomquist et al. 2010). Here, however, we consider the situation where genetic data is available only from a single location, which is subject to imports from other locations about which little is known. This situation occurs frequently, for example due to high discrepancies between the sequencing capacities of high and low income countries. Furthermore, even if limited sequences are available from other locations, biases in sampling between locations can often confuse phylogeographic methods (De Maio et al. 2015).

We therefore address the problem of inferring the number and phylogenetic location of imports into a population based on samples taken only from that population. This problem is important for determining which measures to take in the control of infectious diseases, since different measures are effective against importation and local transmission. It is also important to consider the presence of imports into a population before attempting to reconstruct local transmission chains with one of the recently developed methods for this purpose (Jombart et al. 2014; Didelot et al. 2017; Klinkenberg et al. 2017; De Maio et al. 2016). Only one of these methods considered the possibility of importation by performing a test based on the number of mutations between a case and its most likely donor (Jombart et al. 2014).

Our starting point is a dated phylogeny for the samples at the location of interest. Such a phylogeny can be constructed either directly from the genomes using BEAST (Suchard et al. 2018) or BEAST2 (Bouckaert et al.2019), or by dating the nodes in a standard phylogeny using treedater (Volz and Frost 2017), TreeTime (Sagulenko et al.2018) or BactDating (Didelot et al. 2018). We consider the leaves of this dated phylogeny in increasing order of sampling dates, asking ourselves for each leaf whether it is likely to to be the result of local transmission or importation from external sources. This chronological approach is important to assess the true number of imports: for example if an import occurred followed by local transmission of the imported variant, the first sample from this variant should be labelled as an import, but subsequent samples from the same variant should not. The approach also lends itself naturally to the online assessment of imports as new cases arise, which is often needed when performing infectious disease epidemiology in real time.

Since we do not have any information about the external sources, and do not want to make any assumptions about them, we build statistical models based on the hypothesis of local transmission, which are fitted using Bayesian methods. When a leaf of the dated phylogeny is found to be a bad fit for this local model, we deduce that an importation is likely to have occurred. Our model is based on the coalescent framework (Kingman 1982; Donnelly and Tavare 1995) and in particular its extension to heterochronous sampling (Drummond et al. 2002, 2003). We also use the version of the coalescent model that accounts for variations in the population size (Griffiths and Tavare 1994; Donnelly and Tavare 1995). We use simulated datasets to show that our approach has an excellent specificity and a good sensitivity for the detection of imports. We also show that our approach can be useful in practice by analysing several recently published real datasets.

## MATERIALS AND METHODS

### Coalescent framework and notations

Let *n* denote the number of tips in a dated phylogeny 𝒢, let s_1:*n*_ denote the dates of the leaves and c_1:(*n−*1)_ denote the dates of the internal nodes. Let A(t) denote the number of lineages at time t in 𝒢. This is easily computed as the number of leaves dated after t minus the number of internal nodes dated after t:

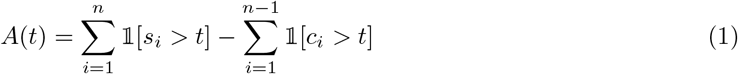

In the coalescent model, each pair of lineages coalesces at rate 1/*N*_e_(t) where *N*_e_(t) is the effective population size at time *t* (Griffiths and Tavare 1994). Note that here and throughout this paper we use the notation *N*_e_ and the name effective population size to denote what is in fact the product of the generation duration and the population size in an idealised Wright−Fisher population. Let us initially assume that this function *N*_e_(*t*) is known, and we will see later how to extend to the situation where it is not known. The total coalescent rate for all pairs at time *t* is therefore equal to:

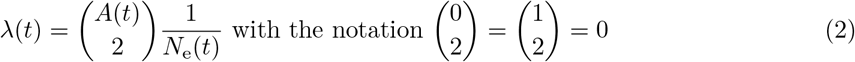

However, here we consider an alternative equivalent formulation of the coalescent model, in which the phylogeny is formed by iterating over the leaves one by one in increasing order of date, and considering how each leaf coalesces with the phylogeny made by the previous leaves (Didelot et al. 2014; Carson et al. 2022). To do so, we consider that the dates *s*_1:*n*_ of the leaves are in increasing order, and that the dates *c*_1:(*n−*1)_ of the internal nodes are ordered so that *c*_*k*_ corresponds to the internal node created when adding the leaf *s*_*k*+1_ to the tree made of the first *k* leaves. Figure 1 shows an example of this notation used for labelling the leaves and nodes of the tree. With these notations, the tree made of the *k* first leaves contains the leaves *s*_1:*k*_ and the nodes *c*_1:(*k−*1)_. We can therefore define the number *A*^*k*^(*t*) of lineages at time t in the tree made of only the first k samples in a way similar to Equation 1:

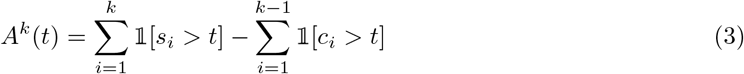

Note in particular that *A*(*t*) from Equation 1 is equal to *A*^*n*^(*t*) from Equation 3 as expected since this corresponds to the number of lineages in the tree made of all *n* leaves and *n −* 1 internal nodes. The rate at which the new leaf s_*k*+1_ coalesces with the tree made of the first *k* leaves is then:

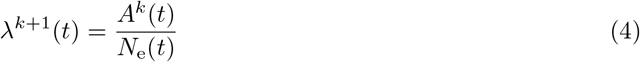

The difference in Equation 4 compared to Equation 2 is that we are now considering coalescence of a single given lineage leading to leaf *s*_*k*+1_, rather than any two pairs of lineages, so that the binomial term for the number of lineages is replaced with simply the number of previous lineages. Note that after the date of leaf *k*, i.e. for *t* > *s*_*k*_, we have no previous lineages, i.e. *A*^*k*^(*t*) = 0, so that λ^*k*+1^(*t*) = 0, i.e. coalescence is impossible. On the other hand, before the date of leaf k, i.e. for *t* < *s*_*k*_, we have always at least one previous lineage so that λ^*k*+1^(t) > 0.

**Figure 1:**
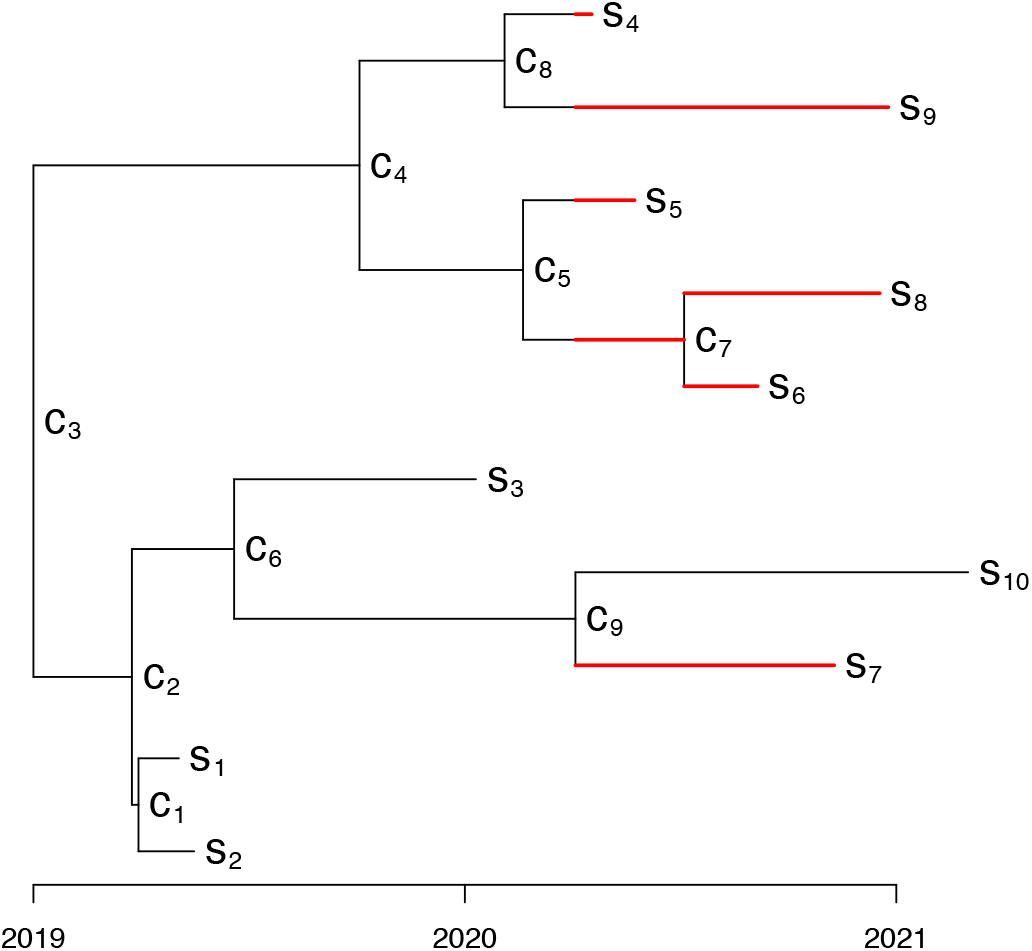
Illustration of the notations used. The leaves are labelled *s*_*k*_ with k in increasing order of sampling date. The internal nodes are labelled c_*k*_ so that leaf *s*_*k*_ coalesces at point *c*_*k−*1_ with the genealogy made of the leaves *s*_1_, …, *s*_*k−*1_. The coalescent interval for leaf s_10_ is shown in red.

Let *C*_*k*_ denote the coalescent interval for leaf *k*, which is defined as the sum of branch lengths between time s_*k*_ and *c*_*k−*1_ in the phylogeny made of the *k −* 1 first leaves. This represents the amount of branch lengths before the new leaf *k* coalesced in the previous tree. Figure 1 shows an example of how the coalescent interval is counted. More formally, we can define the coalescent intervals as:

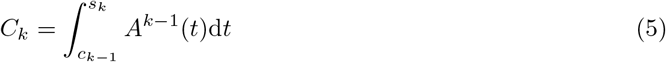

To obtain a given value of *C*_*k*_, we need to have no coalescence of the new lineage between c_*k−*1_ and s_*k*_ and a coalescent event at time c_*k−*1_ with any of the *A*^*k−*1^(*c*_*k−*1_) lineages existing at that time, so that the probability density function of C_*k*_ can be written as:

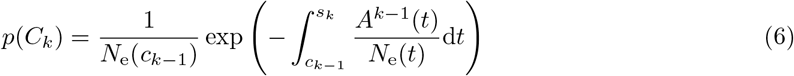

### Detecting imports into a population

Our aim is to find the number and phylogenetic location of import events in a given dated phylogeny. We address this question by considering each leaf of the tree and whether it is likely to be the result of a previously unreported import, given the dated phylogeny made of only the previous samples. If a leaf *s*_*k*_ is not the result of a new import, then its coalescent interval *C*_*k*_ is distributed as described in Equation 6 where *N*_e_(*t*) is the size of the local population. On the other hand, if the leaf *s*_*k*_ is the result of a new import, its coalescent interval will be larger, depending on how distantly related the source of the import is. We do not attempt to explicitly model the source of imports firstly because the data contains little information about import sources, and secondly because we do not want to make assumptions on the sources. We expect most cases in the dated phylogeny to represent local transmission, with only a relatively small ratio (e.g. < 5%) of the number of imports to the number of cases. Only the chronologically first case of any import is classified as such, whereas further cases from the same import represent local transmission following the import.

To progressively explain our methodology for the detection of imports, we will first assume that the demographic function is a known constant, then extend to the case of an unknown constant value, and finally extend to the general case of an unknown variable population size function.

### Case of a known constant population size

Let us first assume that the demographic function *N*_e_(*t*) is a known constant *N*_e_. In this case Equation 6 simplifies into:

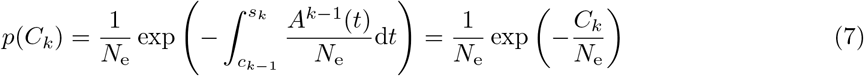

which means that in the case of a constant population size, the coalescent intervals are independent and identically distributed as Exponential with mean *N*_e_. If the leaf *s*_*k*_ is the first reported case of an import, it is likely to have a coalescent interval *C*_*k*_ greater than would be expected if transmission happened only locally, which can be used to form a simple one−sided statistical test with p−value:

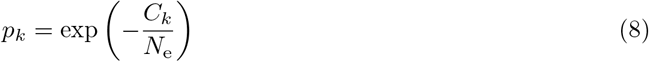

### Case of an unknown constant population size

In the case where the population size function is a constant *N*_e_(*t*) = *N*_e_ which is unknown, we need to estimate it in order to detect imports. We take a Bayesian viewpoint to perform this estimation, which requires setting a prior π(*N*_e_) and combining it with the likelihood terms in Equation 7 to obtain the posterior distribution of *N*_e_:

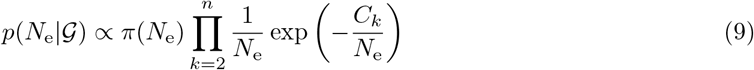

For π(*N*_e_) we use a InvGamma(0.001,0.001) prior, which means that the exponential rate parameter 1/*N*_e_ follows approximately its improper Jeffrey’s prior (Spiegelhalter et al.2002). The Gamma distributions used here and throughout this article are parameterized in terms of the shape and rate parameters, respectively. This same uninformative prior on *N*_e_ was previously used in a method aimed at building dated phylogenies (Didelot et al. 2018).

The posterior distribution in Equation 9 assumes that there are no imports into the local population, so that all coalescent intervals are distributed according to Equation 7. The estimated value of *N*_e_ could therefore be biased upwards compared to the correct value of *N*_e_ in the local population, since any import is likely to have higher coalescent interval values. There are three reasons why this is not a concern in practice. Firstly, we expect only a relatively small number of the leaves to be new imports. Secondly, the distribution of coalescent intervals in the local population (Equation 7) is permissive to high values, so that a few high values do not push up the estimated mean dramatically. Thirdly, if *N*_e_ is overestimated, then we are less likely to detect imports due to having unexpectedly high coalescent intervals. The method is therefore conservative in the detection of imports, rather than having false positives.

We can use a Monte−Carlo approach to generate a sample of *M* values 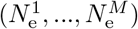 from the posterior distribution in Equation 9, and we can then adapt Equation 8 to compute a posterior predictive p−value (Gelman et al. 1996) to test if the leaf *s*_*k*+1_ is the result of a previously undetected import as:

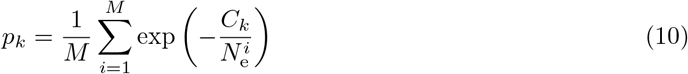

### General case of an unknown

The distribution in Equation 6 represents the model for coalescent intervals if only local transmission occurred and the population size was the function *N*_e_(*t*). However, this equation can not be used in the general case because the demographic function *N*_e_(*t*) is unknown. Phylodynamic methods can be applied to reconstruct the *N*_e_(*t*) function either at the same time as reconstructing a dated phylogeny (Pybus and Rambaut 2009; Ho and Shapiro 2011; Baele et al. 2016) or in a subsequent step (Lan et al. 2015; Karcher et al.2017; Volz and Didelot2018; Didelot et al.2021a). However, all these methods make some assumptions about the demographic function. Furthermore, even if this function was known the resulting distribution for the coalescent intervals in Equation 6 would not be computable analytically. Since our aim here is not to estimate this function but rather to detect imports, we take a different approach.

In the general case, *N*_e_(*t*) is not constant, but with a densely sampled population the coalescent time c_*k−*1_ is likely to be soon before the sampling date *s*_*k*_, so that *N*_e_(*t*) should be approximately constant between c_*k−*1_ and *s*_*k*_ and *C*_*k*_ is approximately exponential as in Equation 7. We therefore consider that the coalescent intervals *C*_*k*_ are exponentially distributed with a mean *µ*(*s*_*k*_) which depends on the date of sampling *s*_*k*_.

To perform Bayesian inference under this model, we need to define the joint prior π(*µ*(s_2_), *µ*(*s*_3_), …, *µ*(*s*_*n*_)). We use a Gaussian process with mean zero and covariance function *k*(*s, s*′) equal to the the Matéern kernel with smoothness *v* = 3/2 (Genton 2002; Williams and Rasmussen 2006):

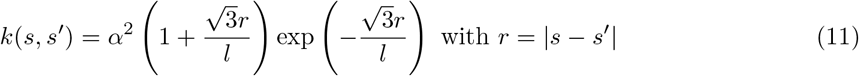

The spectral density function of this kernel in one dimension is:

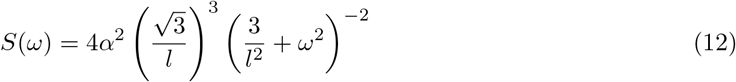

This kernel is characterised by two parameters: the length scale *l* which represents how quickly the distance between two points reduces their correlation, and the scale *α* which represents the marginal standard deviation of the kernel. Specifying the prior on these two parameters completes the definition of the prior model:

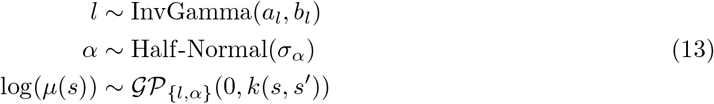

This prior is applied to the dated phylogeny rescaled in the interval [*−* 1, 1], so that the root is at time t = *−* 1 and the most recent leaf at time t = 1. This ensures that the timescale used in the dated phylogeny does not affect the analysis: for example the same dated phylogeny with branch lengths measured in years or in days will produce exactly the same results. In all examples shown here we used hyperparameter values a_*l*_ = b_*l*_ = er_*α*_ = 5. We will show that the choice of these values has little effect on our results.

We want to perform inference in a way that is not computationally intensive even for large phylogenetic trees. Combining this objective with the necessary assumption of dense sampling, together with the assumption that the coalescent rate does not fluctuate too wildly, lends itself naturally to the use of an approximation of the full−rank Gaussian Process. We resort to using a Hilbert space Gaussian process (HSGP) approximation recently described (Riutort−Mayol et al. 2020; Solin and Särkkä 2020). This requires setting two approximation parameters *M* and *L* corresponding to the number of terms in the expansion and the domain size, respectively. We use *M* = 20 and *L* = 2 as previously suggested (Riutort−Mayol et al. 2020).

This model is fitted to the data using the Hamiltonian Monte Carlo method implemented in Stan (Carpenter et al. 2017). For each leaf *s*_*k*_, this results in a Monte−Carlo sample of size *M* denoted (*µ*^1^(*s*_*k*_), …, *µ*^*M*^ (*s*_*k*_)) from the posterior distribution of *µ*(*s*_*k*_). We can then use these values to detect imports using a similar posterior predictive p−value as in Equation 10, namely:

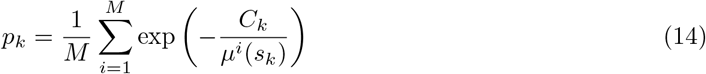

### Implementation

We implemented the simulation and inference methods described in this paper in a new R package entitled *DetectImports* which is available at https://github.com/xavierdidelot/DetectImports. We used the *cmdstanr* package (https://mc-stan.org/cmdstanr/) as interface to Stan (Carpenter et al. 2017) and the *posterior* package (https://mc-stan.org/posterior/) to store and analyse the results. Our default settings (used throughout this article) use 4 chains with 4000 iterations each (1000 for warmup and 3000 for sampling) and an adaptation target acceptance statistic *δ* = 0.9. This number of chains is a choice of convenience, to show that good results can be obtained on a standard laptop, but users have the option to increase this number if wanted. We made sure that no divergent transitions occurred during the sampling phase. Convergence and mixing of the algorithm were verified by checking that for all parameters the improved 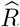 statistics were lower than 1.05 (Vehtarh et al. 2021) and the effective sample sizes greater than 2000. All code and data needed to replicate the results are included in the “run” directory of the *DetectImports* repository.

## RESULTS

### Accounting for variations in the local population size is necessary to correctly identify imports

We can show that the model with constant population size (Equation 10) is insufficient to capture even relatively simple realistic scenarios, and statistical inference based on the variable population size model (Equation 14) is necessary to correctly identify imports. The simulated phylogeny in Figure 2A includes 100 samples taken uniformly throughout a single year, from the 1^st^ January to the 31^st^ December. The ancestral process is the standard coalescent model without any import, but the local effective population size increased five fold in the second half of the year compared to its previous level, from *N*_e_ = 0.2 year to *N*_e_ = 1 year. Consequently the branches tend to be longer in the second half of the year (Figure 2A). We first attempted to detect imports in this phylogeny under our model assuming a constant local population size.This took approximately one second on a standard laptop, and the result is shown in Figure 2B. The mean coalescent interval was estimated to be 0.44 year (with 95% credible interval 0.37−0.54), with the three samples with the largest coalescent intervals having been identified as likely imports (ie. with a posterior predictive p−value *p* < 0.01). This is because these three tips had coalescent intervals higher than would be expected by chance if the population size had been constant, whereas these values were in fact caused by the increase in the population size in the second half of the year. We then inferred using our full model which accounts for variations in the local population size.This took approximately three seconds on a standard laptop, and the results are shown in Figure 2C. The mean coalescent interval was inferred to have increased significantly from the start until the end of 2020, from 0.19 (0.11−0.36) to 1.33 (0.74−2.54). Consequently, the three tips with the largest coalescent intervals were no longer detected as imports, ie the using the full model removed the false positives.

**Figure 2:**
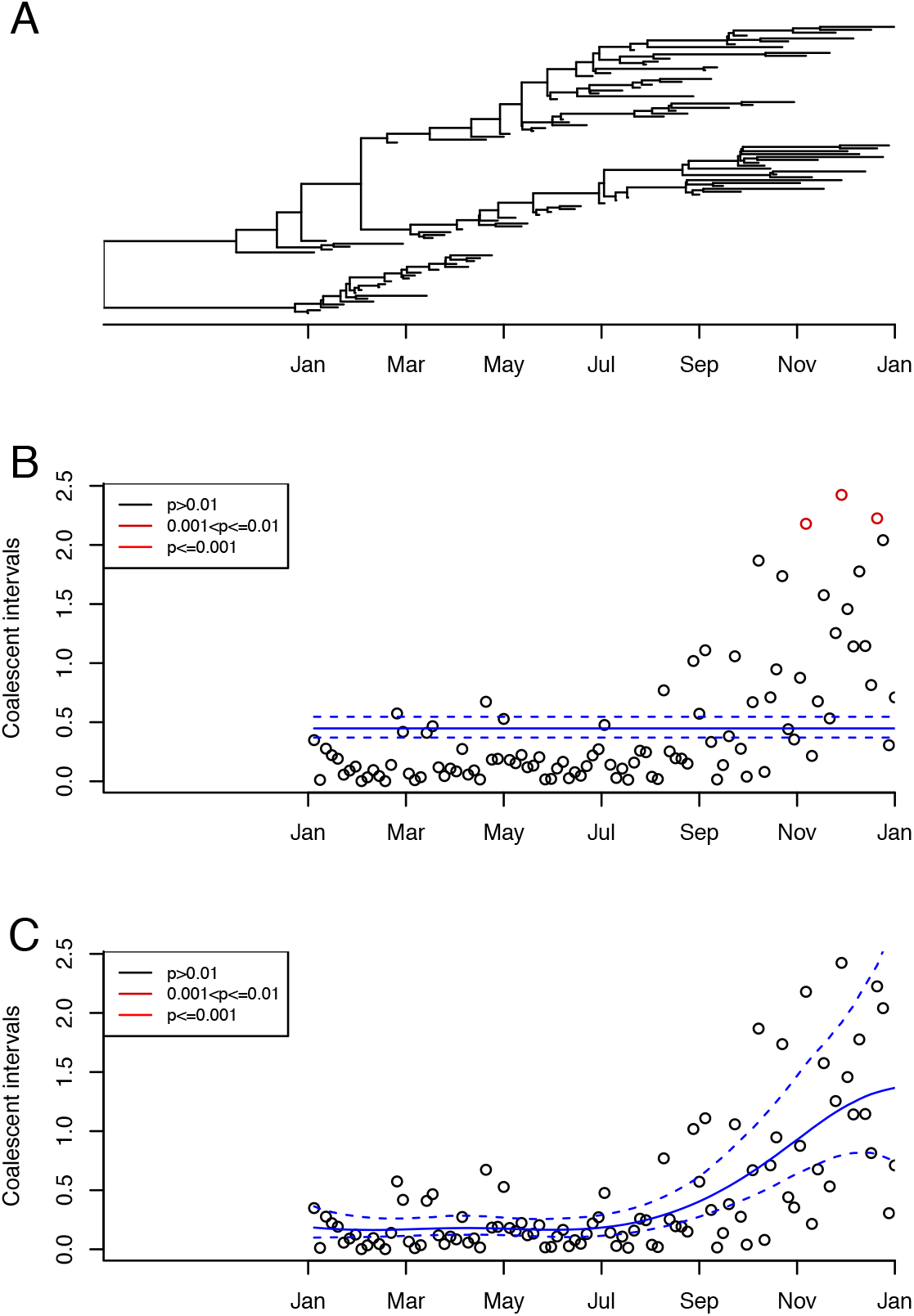
Illustrative application to a single simulated dataset showing that ignoring variations in the local population size can lead to false positives in the detection of imports. A: Simulated phylogeny. B: Inference of imports under the model with constant population size. C: Inference of imports under the model with variable population size. In parts B and C, the inferred mean and 95% credible intervals of the mean coalescent intervals over time are shown in blue.

The example in Figure 2 shows that ignoring the variations in the local population size can lead to the detection of imports that are not real. Conversely, it is important to account for variations in the local population size to avoid real imports going undetected. To illustrate this, we simulated a phylogeny shown in Figure S1A in which the local population starts on 1^st^ January, with a single import happening on 1^st^ April. Both the original and imported strains follow the same linear growth in effective population size *N*_e_(*τ*) = 10*τ* where *τ* is measured in years since the strain introduction. The original and imported strains coalesce together soon before the 1^st^ January. A total of 500 genomes were sampled between the 1^st^ January and the 31^st^ December, with sampling happening at a rate proportional to the effective population size of each strain. When inferring imports under the constant population size model as shown in Figure S1B, the correct import was not detected (*p* = 0.19) but five spurious imports were detected (*p* < 0.01). On the other hand, when inferring imports under the variable population size model as shown in Figure S1C, the correct import was the only one to be detected (*p* = 0.002). The run times were approximately 2 and 13 seconds on a standard laptop computer, for the inference with constant and variable population size, respectively.

We performed one hundred repeats of a similar simulated scenario to the one described above, except that after the local population was initiated on the 1^st^ January, there were two imports in each simulation on the 1^st^ April and on the 1^st^ July. A total of 500 genomes were sampled throughout the year between the 1^st^ January and the 31^st^ December, with sampling happening at a rate proportional to the effective population size of each of the three strains (initial plus two imports). We performed inference under both models with constant and variable population size, and computed the sensitivity and specificity of both import classifiers at different values of the posterior predictive p−values. This resulted in the receiver operating characteristic (ROC) curves shown in Figure 3. The ROC curve for the model with constant population size is far from perfect, with an area under the curve (AUC) of 0.895. This AUC value represents the probability of giving a lower posterior predictive p−value of import to an imported sample compared to a sample that was not imported. In contrast, the model with variable population size has an almost perfect ROC curve, with an AUC of 0.997 (Figure 3). Considering *p* = 0.01 as the cutoff for significance, the inference under the constant population size model has a specificity of 98.6% and a sensitivity of only 40.5%, whereas the inference under the variable size model has a specificity of 99.7% and a sensitivity of 97.0%. To ensure that our choice of the prior did not have undue effect on the results, we repeated this ROC analysis with hyperparameter values a_*l*_ = b_*l*_ = *σ*_*α*_ = 2 and found that it made little difference (Figure S2).

**Figure 3:**
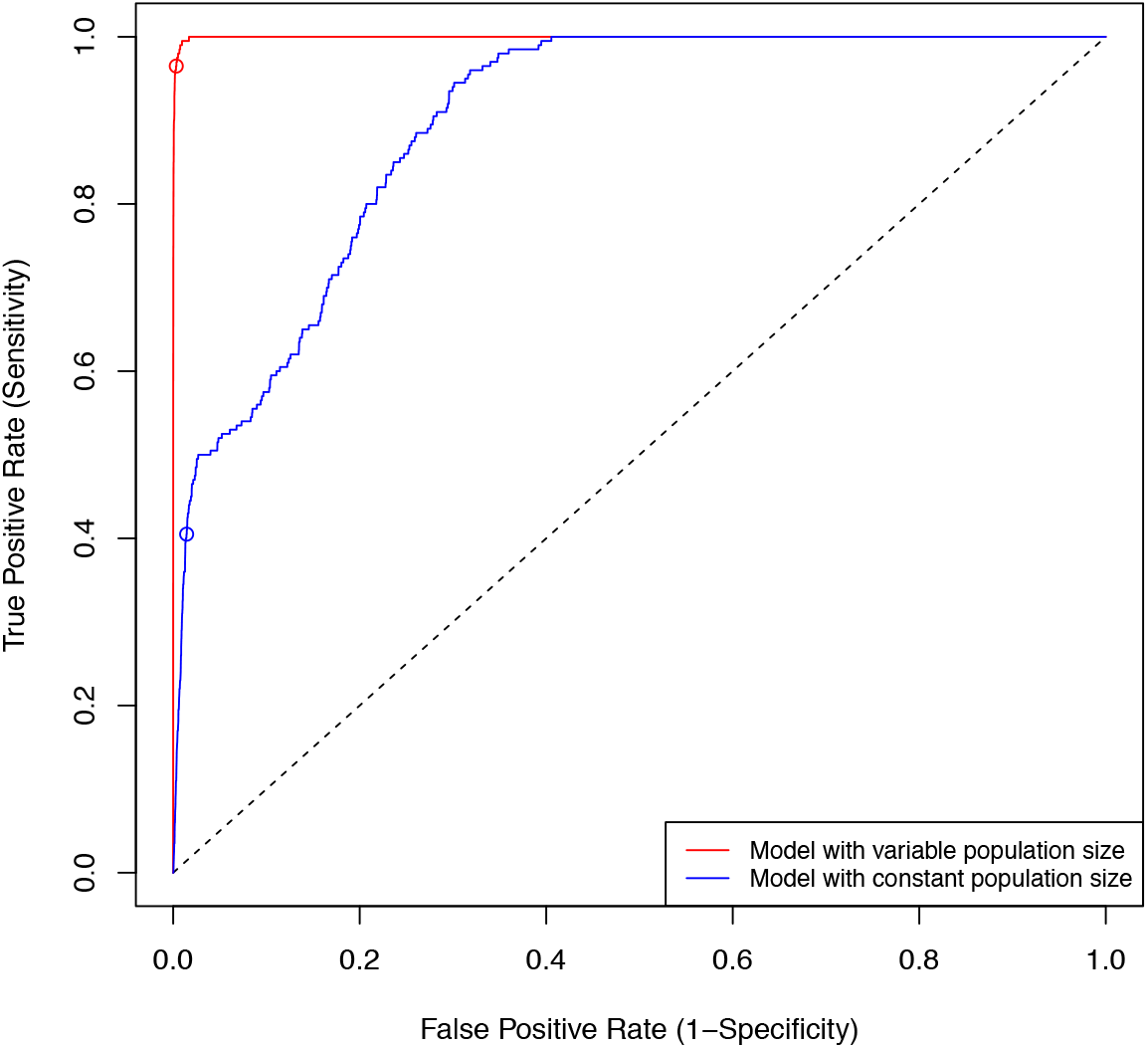
Receiver operating characteristic (ROC) curves for the model with variable population size (red) and the model with constant population size (blue). The dots represent a p−value of 0.01.

This ROC analysis (Figure 3) confirms the result illustrated with specific examples in Figures 2 and S1 about the importance of accounting for the variations in the local population size in order to detect imports with good specificity and sensitivity, and the variable population size model will therefore be used throughout the rest of this paper.

### Inference on simulated datasets from the structured coalescent model

The simulations above were considering only the phylogenetic process within the local population. Here we consider a more complex model in which the global population is structured into several locations, also known as demes, with migrations potentially occurring from any deme to any other. The corresponding genealogical process is described by the structured coalescent model (Notohara 1990; Hudson *1990; Muller et al. 2017). We used the software Master*, a stochastic simulator of birth− death master equations, (Vaughan and Drummond 2013) to simulate under this model with *D* demes which all had the same effective population size *N*_*e*_ = 1 year. The backward−in−time migration rate from any deme to any other was sampled uniformly at random between 0 and 0.5/(*D −* 1), so that the expected waiting time until a migration was the same for all values of D. Only one of the D demes was sampled 500 times with dates taken uniformly at random over a period of a year. We performed 100 simulations with *D* = 5, *D* = 3 and *D* = 2, each.

For each simulated dataset, we counted the correct number of imports into the local population by looking through the whole migration history for migrations into the local deme that led directly (ie without any other migration event on the phylogenetic path) to at least one sampled leaf. We also inferred the number of imports based on the dated phylogeny of the samples from local deme, which took between 15 and 20 seconds to run for each simulated dataset. Figure 4 compares the correct and inferred number of imports in each simulation. The number of detected imports is correlated with the correct number of imports in all three cases with *D* = 5 demes (Figure 4A), *D* = 3 demes (Figure 4B) and *D* = 2 demes (Figure 4C). However, in all three cases we find that the number of imports has been estimated, with on average only 81%, 76% and 69% of imports being detected for *D* = 5, *D* = 3 and *D* = 2 demes, respectively. This increasing relationship between the number of demes and the ability to detect imports into one of the demes is as expected: when the number of demes is larger, the local population represents a smaller proportion of the global population. Each import becomes more clearly separated in the phylogenies and therefore easier to detect. The fact that some imports remain impossible to detect in all three cases is also expected, since there is always the possibility that a lineage going back in time migrates out of the local population and back into it quickly afterwards, making it basically undetectable. Finally, the case with *D* = 2 demes is especially interesting since in this case there are just two populations of equal sizes, one which is sampled and the other one not. Detecting imports is clearly challenging in these conditions, harder than we would envisage in most applications to real data where the local population would typically be a small fraction of the global population. It is therefore encouraging to see that even in this difficult case our method was able to detect the majority of the imports (Figure 4C).

**Figure 4:**
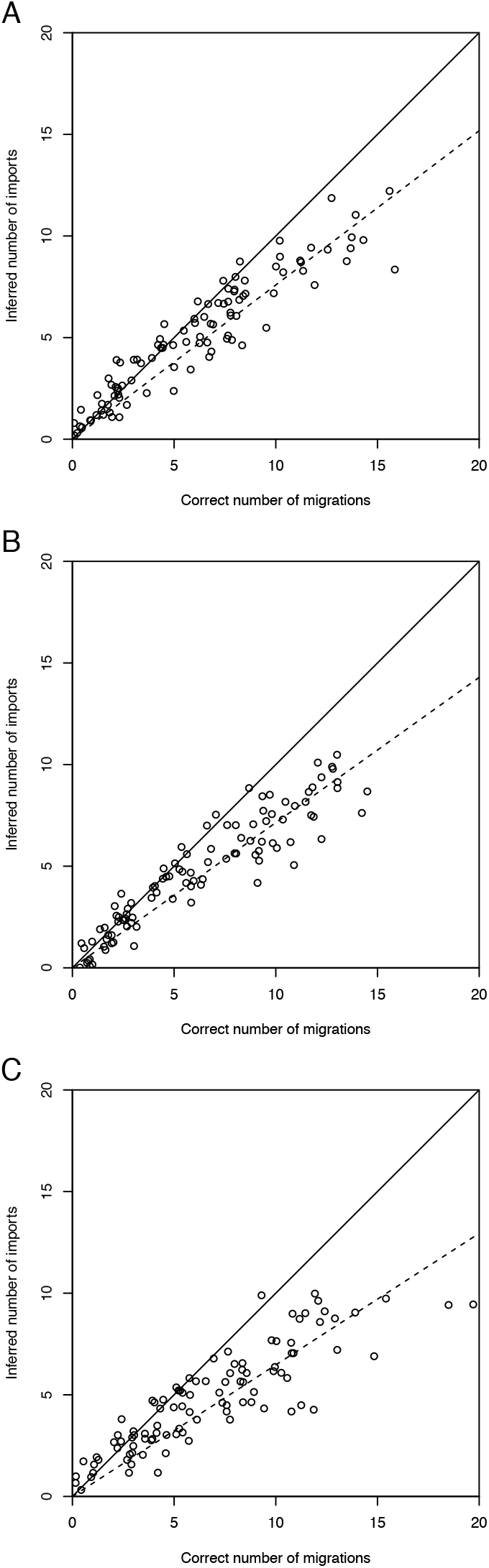
Application to simulated datasets from the structured coalescent model with 5 demes (A), 3 demes (B) and 2 demes (C).

### Application to real datasets

We also considered applications to real datasets, and considered how our inference compares with accepted epidemiological wisdom about a number of outbreaks originated by diverse pathogens.

First we analysed a small dataset of 132 genomes from an outbreak of *Neisseria gonorrhoeae* (Didelot et al. 2016). The genomes were collected between 1995 and 2000 as part of a prospective study on gonorrhoea in Sheffield (Ward et al. 2000), and all belonged to ST12 which was the most prevalent NG−MAST type in this setting (Bilek et al. 2007). In the previous study of this data (Didelot et al. 2016), a dated phylogeny was built using BEAST (Suchard et al. 2018) as shown in Figure S3A. Analysis took approximately 5 seconds and the result is shown in Figure S3B. No import was found in this dataset, confirming that all the genomes belonged to the same outbreak and can be analysed as such as previously performed (Didelot et al. 2016). In particular, there was a gap of about 2 years in the sampling in 1998 and 1999, with most genomes originating before this gap and only seven genomes corresponding to cases afterwards. In principle, this gap could have been explained by a clearance and reintroduction of ST12 in the region, but our analysis shows that this is not the case. Instead, the later cases are descended from the earlier ones through chains of unsampled transmission intermediates, as previously proposed (Didelot et al. 2016) using *outbreaker* (Jombart et al. 2014).

Second, we analysed a collection of 155 Vietnamese genomes from the VN clade of the emerging enteric pathogen *Shigella sonnei* (Holt et al. 2013). These genomes were sampled between 1995 and 2010, and a dated phylogeny was built using the additive relaxed clock model in BactDating (Didelot et al. 2021b). The import analysis took approximately 6 seconds and the result is shown in Figure 5. A single import was found (isolate labelled 30451) with a posterior predictive p−value of 0.0057. This isolate may not look remarkably different at first sight on the phylogeny (Figure 5A), but it is the second most recent isolate in the collection and has by far the largest coalescent interval (Figure 5B). We repeated this analysis for 100 phylogenies from the posterior sample produced by BactDating (Figure S4). The results were robust to phylogenetic uncertainty, with only isolate 30451 being a likely import. The p−values for this isolate had an interquantile range between 0.007 and 0.014 (Figure S4). Given these p−values we can not be absolutely certain if this isolate was indeed imported, but if so it would most probably represent a relatively quick migration out and back into the Vietnamese population, for example via a neighbouring country.

**Figure 5:**
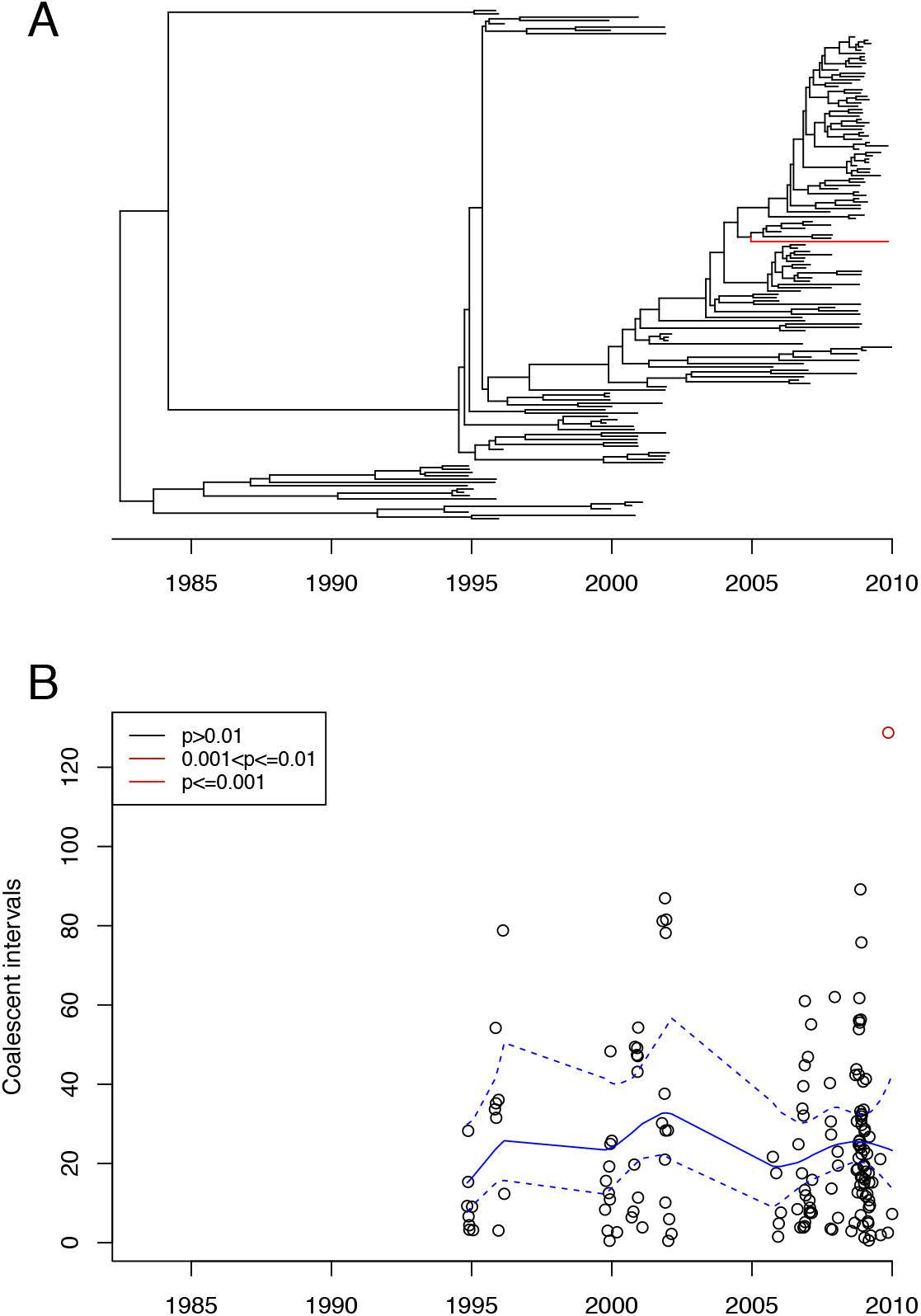
Application to *Shigella sonnei* dataset. A: Dated phylogeny with imports highlighted in red. B: Inference of imports. The inferred mean and 95% credible intervals of the mean coalescent intervals over time are shown in blue.

We also analysed a collection of 1031 genomes of Ebola isolated from Sierra Leone between the 25^th^ May 2014 and 12^th^ September 2015. A dated phylogeny was built for these genomes using BEAST (Suchard et al. 2018) in a previous study (Dudas et al. 2017). The import analysis took 50 seconds. The results are shown in Figure 6, with a total of 26 imports found. All the inferred imports correspond to isolates from 2015, despite most (544/1031) isolates in this collection being from 2014, which is statistically significant (Fisher’s exact test, *p* < 10^*−*4^). This result coincides well with the incidence of Ebola over time in Sierra Leone and the two other badly affected neighbouring countries Guinea and Liberia (Shultz et al.2016). The end of 2014 and beginning of 2015 corresponds to the time when Sierra Leone managed to greatly reduce the number of Ebola cases, whereas other countries took longer to do so. It is also interesting to compare our results with a phylogeographic study of genomes from Sierra Leone, Guinea and Liberia (Dudas et al. 2017). This study found that the vast majority of cases in Sierra Leone were descended from an initial introduction of Ebola from Guinea in early 2014, but that a few sporadic cases were linked with several reintroduction events from Guinea dated between January and April 2015, which is in very good agreement with our results based on genomic data from Sierra Leone only (Figure 6).

**Figure 6:**
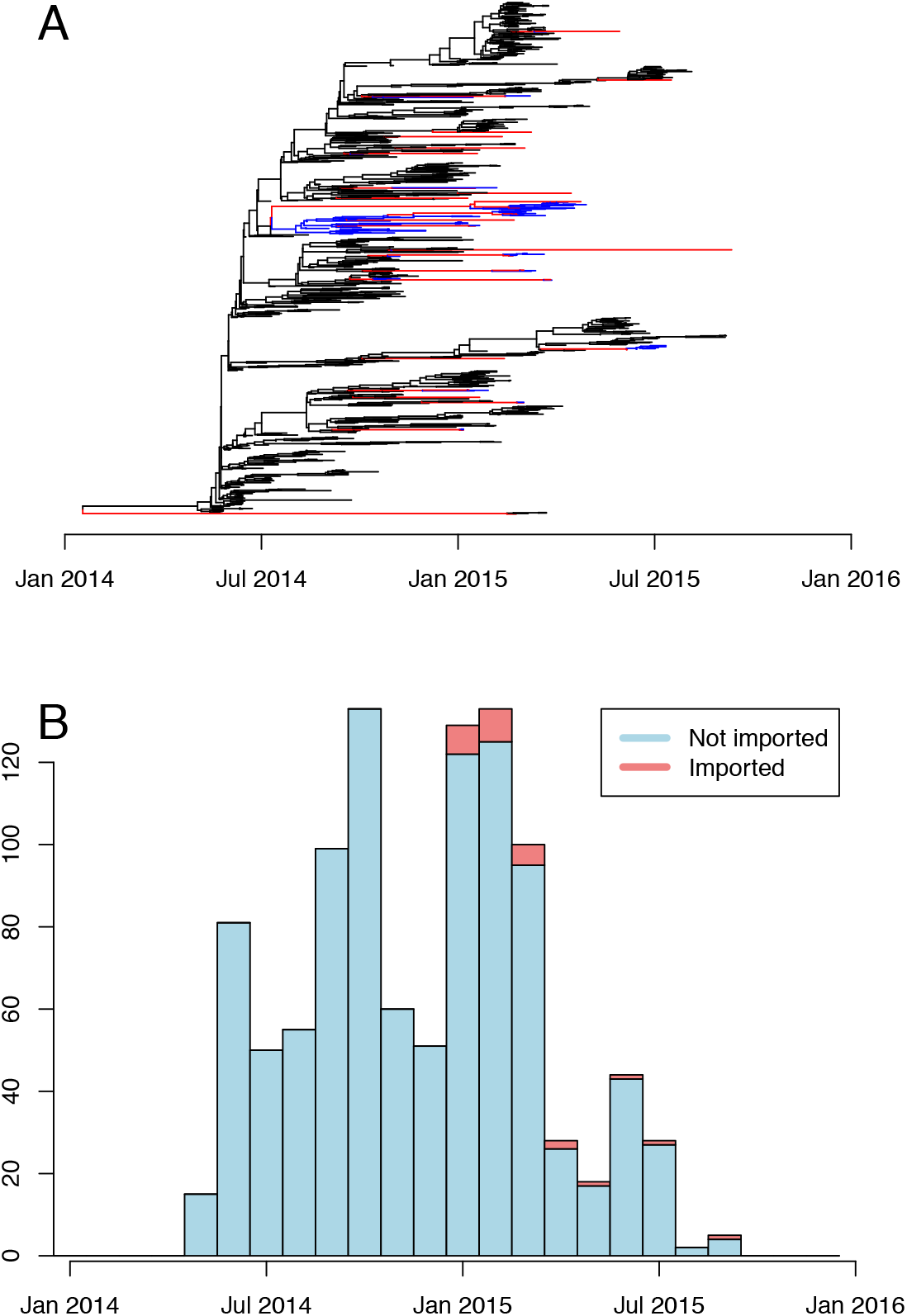
Application to Ebola dataset. A: Dated phylogeny showing the imports (red) and locally transmitted descendants of imports (blue). B: Histogram showing the number of locally transmitted (blue) and imported (red) isolates over time.

Finally, we analysed a set of 3797 SARS−CoV−2 genomes isolated in Scotland between March 2020 and June 2022. This collection was obtained by downsampling the ∼200,000 Scottish sequences that have been deposited in GISAID since the beginning of the COVID−19 pandemic; for each day for which data was available, at most 5 genomes were randomly selected from those having no ambiguous or unknown base. We then cleaned up the multiple sequence alignment retrieved from GISAID by only keeping the relevant rows, eliminating columns entirely made of dashes, and trimming sequences at both sides by the minimum amount of nucleotides needed to make the stretches of dashes at the beginning and the end of each genome, which indicate unknown sequence, entirely disappear for all the genomes selected. The resulting alignment was given as input to FastTree (Price et al. 2010) to generate a phylogeny which was then dated using BactDating (Didelot et al.2018). The inference of imports took approximately 20 minutes to compute, resulting in a total of 50 detected imports, as shown in Figure 7 and listed in Table S1. Interestingly, no imports were found until August 2020, perhaps as an effect of the first lockdown which started at the end of March 2020 and was progressively relaxed throughout Spring 2020. During August 2020 15 imports were identified, which was the largest for any month in the analysis. Many of these imports may be associated with Summer holidaying. According to our analysis the alpha variant was imported in November 2020, soon after it had been reported in England (Davies et al. 2021). Several imports corresponded to low frequency variants, including Beta in December 2020, Zeta in December 2020 and Eta in March 2021. Fewer imports were detected in the first few months of 2021, which may be the result of the second lockdown in January and February 2021. The Delta variant was imported in April 2021, soon before it became dominant throughout the UK (Elliott et al. 2021). From then on, the Alpha variant was reimported three times and the Delta variant nine times with the last imports occurring in December 2021. The Omicron variant (Cao et al. 2022) was first imported in December 2021, and reimported three times from January to March 2022, by which time this variant had become dominant in the UK and globally.

**Figure 7:**
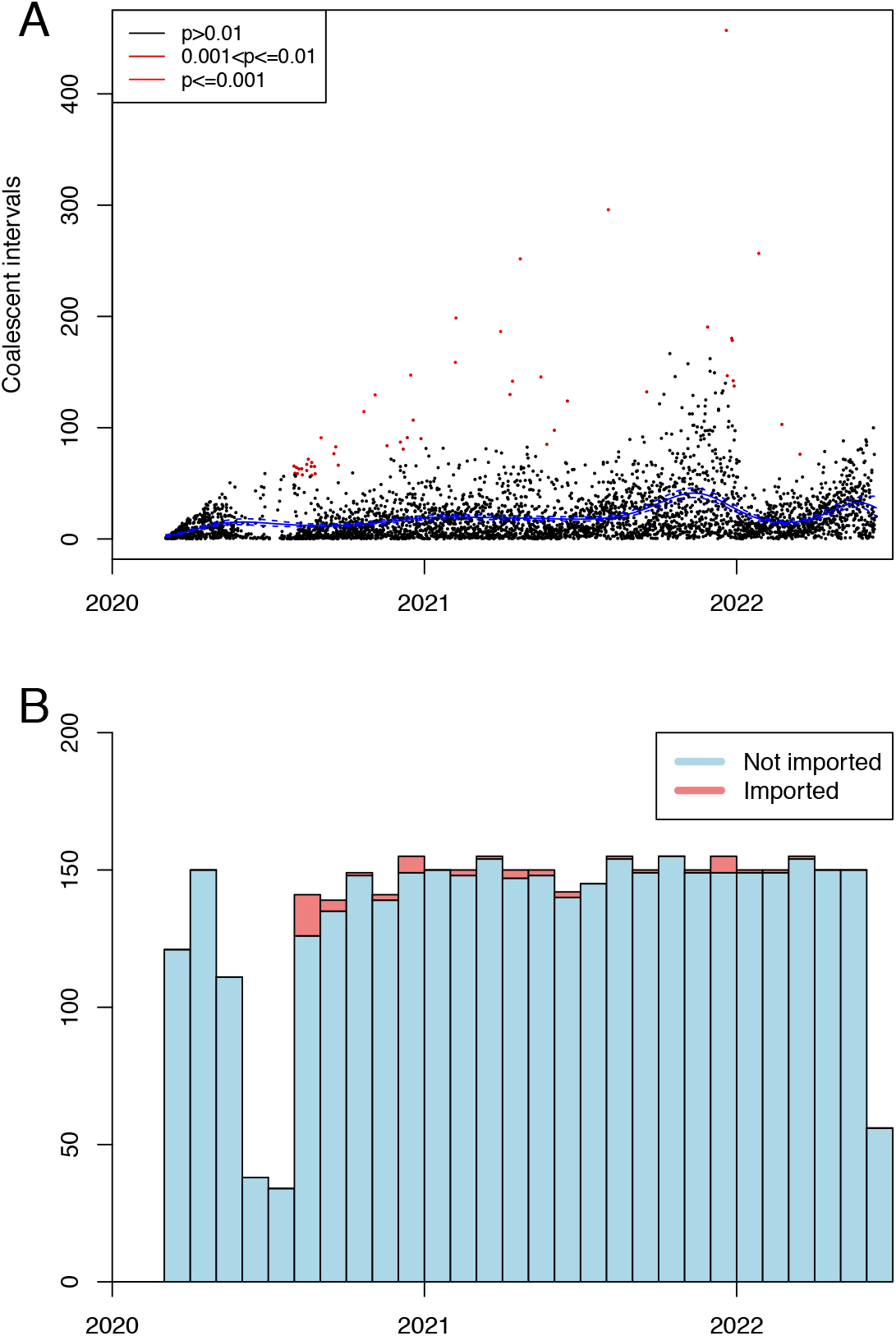
Application to SARS−CoV−2 dataset. A: Inference of imports. The inferred mean and 95% credible intervals of the mean coalescent intervals over time are shown in blue. B: Histogram showing the number of locally transmitted (blue) and imported (red) isolates over time.

## DISCUSSION

When studying the occurrence of an infectious disease in a geographically limited population, it is often important to distinguish cases that have been transmitted within the population from cases that have been imported from external origins. Genomic data has the potential to distinguish between these two types of cases, since new imported cases would usually be more distantly related from previous cases than cases arising from local transmission. We developed a statistical method that can quickly establish which cases have been imported. Application to simulated datasets showed that our method has excellent specificity, which means a very low probability that a locally transmitted case would be inferred to have been imported. Our method also has good sensitivity to detect cases that have been truly imported, although this is not perfect since there is always a chance that an import will be genetically similar to the locally transmitting population. We also showed that our method can be useful in four very different real applications: an outbreak of gonorrhoea in a single city, a country−wide expansion of a bacterial clone causing enteric disease, the 2013−2016 epidemic of Ebola virus disease in Sierra Leone, and the COVID−19 pandemic.

Our approach uses only genomic data from within the location of interest, without making assumptions about the genomic epidemiology of the disease outside of this location. This problem is therefore analogous to the inference of recombination coming from external unsampled sources (Didelot and Falush2007; Didelot and Wilson 2015) rather than recombination within a single population (Didelot et al.2010). In this case, genomes from other populations are sometimes used subsequently, by comparing them with the inferred recombination tracts to determine if they might be the origin of the recombination events (Didelot et al. 2011, 2012; Ozer et al. 2019). In the same way, for the problem of detecting imports into a location we are interested in here, any information about the genetic diversity at other locations could be used to assess the likely origin of the detected imports, simply by comparing the genomic sequences of the inferred imports with the genomes collected from other locations.

Our method requires first to compute a dated phylogeny from the genomes before detecting imports, and therefore fits within the framework of step−by−step approaches from microbial genomes to epidemiology (Didelot and Parkhill 2022). There are several advantages to this type of approach, including scalability to large datasets as we demonstrated here with the analysis in a matter of seconds of datasets containing hundreds of pathogen genomes. There are however also drawbacks to such an approach compared to a more integrated approach (Didelot and Parkhill 2022). A first issue concerns the fact that the model used to build the dated phylogeny is contradicted here by the presence of imports. As previously noted (cf Methods), this is unlikely to have a significant effect as long as imports are relatively rare, but in any case the effect would be to overestimate the local effective population size, thus making the method more specific and less sensitive, as desired. Another inaccuracy of the step−by−step approach is that a single dated phylogeny is used as input, which does not capture the uncertainty in the phylogenetic reconstruction. A solution is to apply the method to a posterior sample of the dated phylogenies (Nylander et al. 2008), which is feasible here since our method to detect imports is very fast. We applied this idea to one of the real datasets we analysed and found that the detection of imports was relatively robust even when using a single consensus tree (Figure S4). This is as expected since imports correspond to long branches of the tree which are unlikely to have much uncertainty.

## ACKNOWLEDGEMENTS

We acknowledge funding from the National Institute for Health Research (NIHR) Health Protection Research Unit in Genomics and Enabling Data. This work was supported by the UK Engineering and Physical Sciences Research Council (EPSRC) grant EP/S022244/1 for the EPSRC Centre for Doctoral Training in Mathematics for Real−World Systems II. XD and PR acknowledge the Research/Scientific Computing teams at The James Hutton Institute and NIAB for providing computational resources and technical support for the UK’s Crop Diversity Bioinformatics HPC (BBSRC grant BB/S019669/1), use of which has contributed to the results reported within this paper.

