## Supplementary Material for "Distinguishing imported cases from locally acquired cases within a geographically limited genomic sample of an infectious disease"

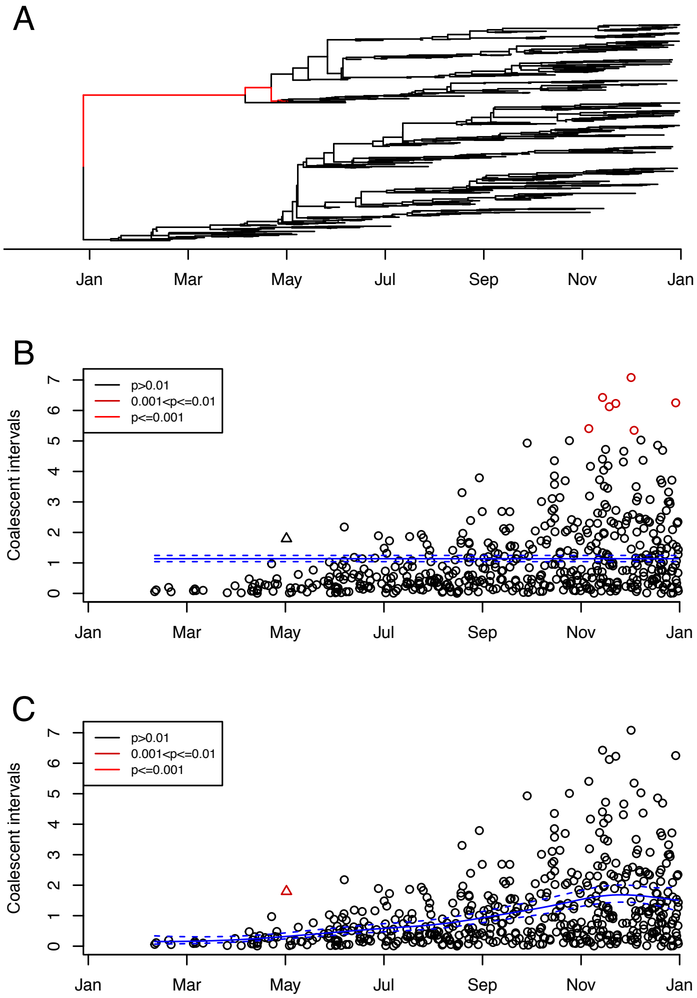


Figure S1: Illustrative application to a single simulated dataset showing that ignoring variations in the local population size can lead to false negatives in the detection of imports. A: Simulated phylogeny. B: Inference of imports under the model with constant population size. C: Inference of imports under the model with variable population size. In parts B and C, the inferred mean and 95% credible intervals of the mean coalescent intervals over time are shown in blue. The triangle in parts B and C corresponds to the first imported case shown in red in part A.

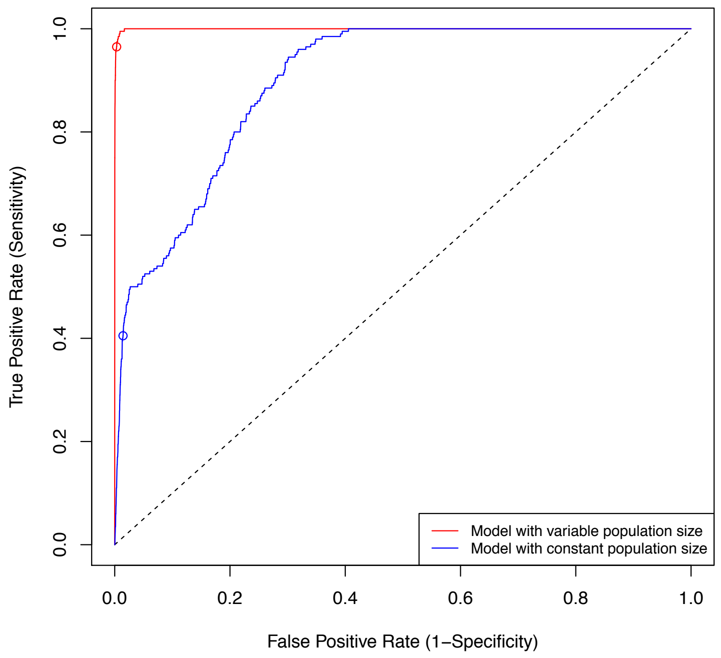


Figure S2: Receiver operating characteristic (ROC) curves for the model with variable population size (red) and the model with constant population size (blue). The dots represent a p-value of 0.01. Hyperparameter values $a_l = b_l = \sigma_\alpha = 2$ were used instead of $a_l = b_l = \sigma_\alpha = 5$ as in Figure 3.

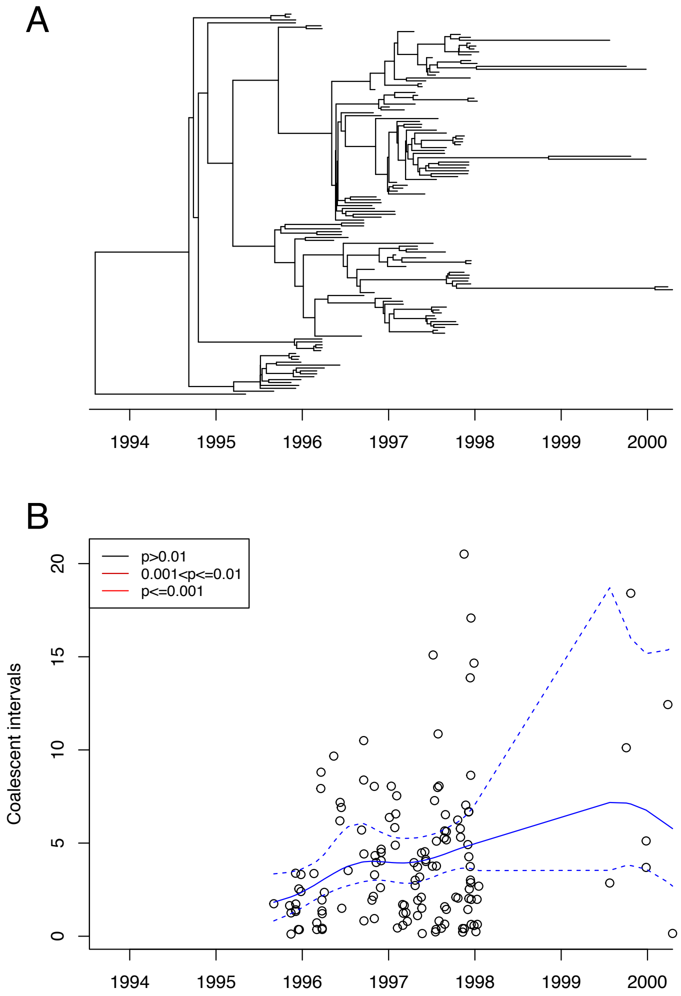


Figure S3: Application to an outbreak of gonorrhoea. A: Dated phylogeny. B: Inference of imports. The inferred mean and 95% credible intervals of the mean coalescent intervals over time are shown in blue.

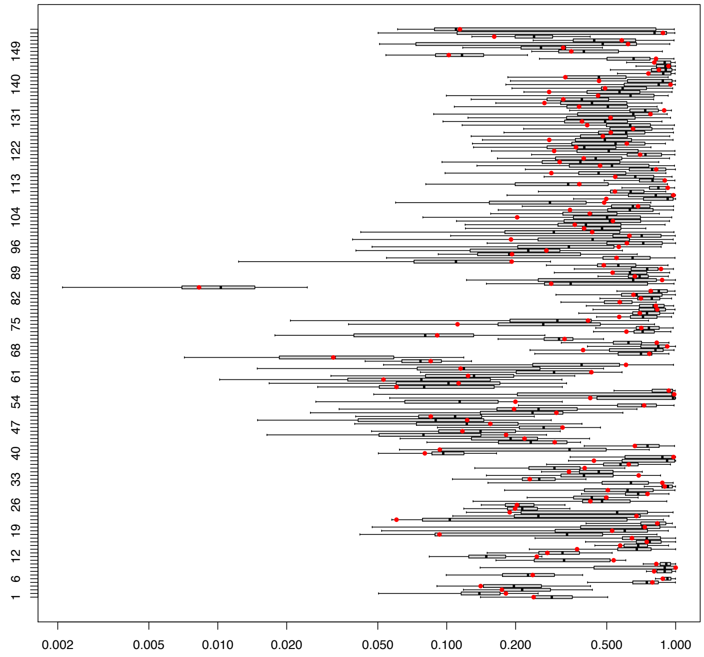


Figure S4: Application to *Shigella sonnei* dataset based on 100 dated phylogenies from the BactDating posterior sample. The boxplots show the distributions of p-value obtained for each case, and the red dots show the p-value computed from the consensus phylogeny.

| Genome | Date | Lineage | Variant |
|---|---|---|---|
| EPI_ISL_532587 | 8/1/20 | AD.2 | |
| EPI_ISL_532620 | 8/2/20 | B.1.1 | |
| EPI_ISL_532118 | 8/4/20 | B.1.389 | |
| EPI_ISL_532024 | 8/5/20 | B.1.235 | |
| EPI_ISL_532105 | 8/7/20 | B.1.1.279 | |
| EPI_ISL_534413 | 8/10/20 | B.1.1.372 | |
| EPI_ISL_531473 | 8/11/20 | B.1.1.311 | |
| EPI_ISL_534647 | 8/16/20 | B.1.1.303 | |
| EPI_ISL_568421 | 8/16/20 | B.1.258 | |
| EPI_ISL_568423 | 8/18/20 | B.1.441 | |
| EPI_ISL_530768 | 8/21/20 | B.1.221 | |
| EPI_ISL_529685 | 8/22/20 | B.1.282 | |
| EPI_ISL_568393 | 8/22/20 | B.1.416 | |
| EPI_ISL_568293 | 8/25/20 | B.1.1 | |
| EPI_ISL_568360 | 8/26/20 | B.1.1.277 | |
| EPI_ISL_577371 | 9/2/20 | B.1.1 | |
| EPI_ISL_601940 | 9/17/20 | B.1.1.255 | |
| EPI_ISL_585313 | 9/19/20 | B.1 | |
| EPI_ISL_601175 | 9/22/20 | B.1.1.307 | |
| EPI_ISL_623930 | 10/22/20 | B.1.1.170 | |
| EPI_ISL_676414 | 11/4/20 | B.1.1.7 | Alpha |
| EPI_ISL_757190 | 11/18/20 | B.1.1.317 | |
| EPI_ISL_710633 | 12/4/20 | B.1.36.26 | |
| EPI_ISL_757231 | 12/7/20 | B.1.36.29 | |
| EPI_ISL_919311 | 12/12/20 | B.1.177.17 | |
| EPI_ISL_989986 | 12/16/20 | B.1.351 | Beta |
| EPI_ISL_997124 | 12/19/20 | P.2 | Zeta |
| EPI_ISL_865085 | 12/28/20 | B.1.177 | |
| EPI_ISL_1386678 | 2/6/21 | B.1.562 | |
| EPI_ISL_1247731 | 2/7/21 | B.1.1.318 | |
| EPI_ISL_1538076 | 3/31/21 | B.1.525 | Eta |
| EPI_ISL_1673240 | 4/11/21 | B.1.1.7 | Alpha |
| EPI_ISL_1741634 | 4/14/21 | B.1.1.7 | Alpha |
| EPI_ISL_1829569 | 4/23/21 | B.1.617.2 | Delta |
| EPI_ISL_2236724 | 5/17/21 | B.1.617.2 | Delta |
| EPI_ISL_2395821 | 5/24/21 | B.1.617.2 | Delta |
| EPI_ISL_2487803 | 6/2/21 | B.1.1.7 | Alpha |
| EPI_ISL_2700963 | 6/17/21 | B.1.1.7 | Alpha |
| EPI_ISL_3529997 | 8/4/21 | P.1 | Gamma |
| EPI_ISL_4600095 | 9/18/21 | AY.34 | Delta |
| EPI_ISL_7306019 | 11/28/21 | AY.109 | Delta |
| EPI_ISL_8763944 | 12/20/21 | BA.1 | Omicron |
| EPI_ISL_8197175 | 12/21/21 | AY.120 | Delta |
| EPI_ISL_8457600 | 12/26/21 | AY.4 | Delta |
| EPI_ISL_8457302 | 12/27/21 | AY.98 | Delta |
| EPI_ISL_8456419 | 12/28/21 | AY.4 | Delta |
| EPI_ISL_8440261 | 12/29/21 | AY.4 | Delta |
| EPI_ISL_9510674 | 1/27/22 | BA.2.10 | Omicron |
| EPI_ISL_10641104 | 2/23/22 | XE | Omicron |
| EPI_ISL_11401173 | 3/16/22 | BA.2 | Omicron:1 |

Table S1: List of imports found in the SARS-CoV-2 dataset.
